# In Situ-Crosslinked Zippersomes Enhance Cardiac Repair by Increasing Accumulation and Retention

**DOI:** 10.1101/2024.03.14.585030

**Authors:** Natalie E. Jasiewicz, Kuo-Ching Mei, Hannah M. Oh, Emily E. Bonacquisti, Ameya Chaudhari, Camryn Byrum, Brian C. Jensen, Juliane Nguyen

## Abstract

Mesenchymal stem cell (MSC)-derived extracellular vesicles (EVs) are a promising treatment for myocardial infarction, but their therapeutic efficacy is limited by inefficient accumulation at the target site. A non-invasive MSC EV therapy that enhances EV accumulation at the disease site and extends EV retention could significantly improve post-infarct cardiac regeneration. Here we show that EVs decorated with the next-generation of high-affinity heterodimerizing leucine zippers, termed high-affinity (HiA) Zippersomes, amplify targetable surface areas through in situ crosslinking and exhibited ∼7-fold enhanced accumulation within the infarcted myocardium in mice after three days and continued to be retained up to day 21, surpassing the performance of unmodified EVs. After myocardial infarction in mice, high-affinity Zippersomes increase the ejection fraction by 53% and 100% compared with unmodified EVs and PBS, respectively. This notable improvement in cardiac function played a crucial role in restoring healthy heart performance. High-affinity Zippersomes also robustly decrease infarct size by 52% and 60% compared with unmodified EVs and PBS, respectively, thus representing a promising platform for non-invasive vesicle delivery to the infarcted heart.

**Translational Impact Statement:** Therapeutic delivery to the heart remains inefficient and poses a bottleneck in modern drug delivery. Surgical application and intramyocardial injection of therapeutics carry high risks for most heart attack patients. To address these limitations, we have developed a non-invasive strategy for efficient cardiac accumulation of therapeutics using in situ crosslinking. Our approach achieves high cardiac deposition of therapeutics without invasive intramyocardial injections. Patients admitted with myocardial infarction typically receive intravenous access, which would allow painless administration of Zippersomes alongside standard of care.

## Introduction

Despite the widespread use of stem cells for the treatment of myocardial infarction (MI), there is a consensus within the field that acellular approaches may pose fewer barriers to clinical translation and therefore should be considered as therapeutic options. Extracellular vesicles (EVs) – lipid vesicles secreted by cells – provide a middle ground between cellular and synthetic approaches. EVs contain many of the components typically found in their parent cells including proteins, RNAs, growth factors, and chemokines [1–4] but, in contrast to cells, are easier to handle, store, and ship. Their small size, physicochemical properties, and low immunogenicity permit deep tissue penetration, facilitate intercellular communication, and mediate tissue regeneration [5–11]. While many studies have directly injected EVs into the myocardium to deliver them to the target site, this approach is limited by rapid EV elimination, requires open chest surgery, can cause arrhythmias, and can have deleterious effects on cardiac function [12]. Other therapies attempt to circumvent this shortcoming by administering EVs systemically; however, rapid washout and poor accumulation limit their efficacy [13]. Therefore, approaches that can increase EV exposure within the infarct site, preferably non-invasively, are desperately needed.

Inefficient delivery to the infarct site is an issue not limited to EVs but other nanocarriers as well. Though widely used, nanoparticles often suffer from poor homing and rapid, non-specific biodistribution [14]. For example, micelles and liposomes can accumulate in the infarcted myocardium for approximately one day but cannot be detected long-term [15]. Lundy et al. demonstrated that less than 0.5% of systemically administered nanoparticles (20 nm - 2 µm) are retained after 30 minutes[16]. Other studies have shown that nanoparticles that serve as drug carriers can prolong drug retention, but not long-term. For example, PLGA nanoparticles loaded with IGF-1 prolonged delivery for at least 24 hours but not more than three days [17]. Many strategies to enhance the accumulation of nanocarriers or EVs at the infarct site focus on improving nanoparticle targeting or use of additional nanomaterials, such as hydrogels and nanocomplexes[18]. However, these approaches can be limited by low loading efficiency and the need for additional materials. Therefore, a simple, efficient strategy at the nanoscale level would be highly desirable to significantly advance the field.

We previously showed that mesenchymal stem cells (MSCs) decorated with either **b**ase **l**eucine **z**ippers (B-LZ) or their heterodimerizing, **c**omplementary **l**eucine **z**ippers (C-LZ), can crosslink *in situ* to form a cellular depot at the infarct site[19]. Characteristically, leucine zippers are composed of heptad sequences, with leucines every seven residues [20]. Their stable but reversible binding is facilitated by a combination of hydrophobic, hydrophilic, and electrostatic interactions [21–25]. To improve the utility of our protein-based system, we hypothesized that strengthening leucine zipper binding by improving the positions of each intermolecular interaction within the heterodimer in our newly optimized high-affinity leucine zippers would (i) enhance vesicle cross-linking; and (ii) *in vivo*, under systemic flow conditions, better resist venous washout and promote cellular internalization.[20][21–25] By improving the positions of each of these interactions, we sought to increase intermolecular interactions within the dimer and improve binding strength. To enable precise crosslinking and enhance the targetable surface areas, we employ the strategic utilization of heterodimeric leucine zippers exclusively, rather than homodimers. This deliberate choice ensures controlled capturing of each subsequent dose of EVs by every administered dose, thereby amplifying the targetable surface area for enhanced accumulation and retention. This intentional approach effectively safeguards against unintended aggregation and maintains optimal functionality.

Here, we report the development of the next-generation delivery platform of therapeutics to the infarcted myocardium. We provide the first evidence for a non-invasive, in situ crosslinking of small EVs to mediate enhanced and prolonged retention in the infarcted myocardium. We further assess the effect of a newly designed, high-binding affinity leucine zipper on the in vivo crosslinked drug depot. To achieve this, we first decorated the EVs with leucine zippers, termed Zippersomes, and then assessed their ability to crosslink, heterospecifically, *in vitro*. Next, we examined whether the high-binding affinity leucine zippers enhanced EV accumulation and retention in a murine model of MI compared to unmodified EVs and EVs modified with the first-generation of leucine zippers. Finally, we tested the effect of Zippersome treatment on cardiac function and infarct size.

## Results

### Discovery of high-affinity leucine zippers

To enhance the accumulation and retention of intravenously injected EVs for the treatment of MI, we sought to create a second-generation approach to our ZipperCell platform with small nanosized EVs [39]. By utilizing heterodimerizing leucine zippers and decorating them on vesicle surfaces, we aimed to facilitate vehicle crosslinking, in situ. To further improve accumulation in vivo, we created a small library of mutated versions of complementary leucine zipper, C-LZ 10nM to increase binding affinity to the base leucine zipper (B-LZ). By taking the hydrophobic, hydrophilic, and electrostatic interactions that reinforce and stabilize leucine zipper dimers into account (**Fig. 1A**), we carefully selected a group of mutations for our study; three non-interacting surface (NIS) mutations, three interfacial contact (IC) mutations, and three sequences that were a combination of NIS and IC mutations (**Fig. 1B**). The binding affinities of these nine sequences were first determined through molecular docking simulations and then rank ordered based on their binding strengths.

**Figure 1.**
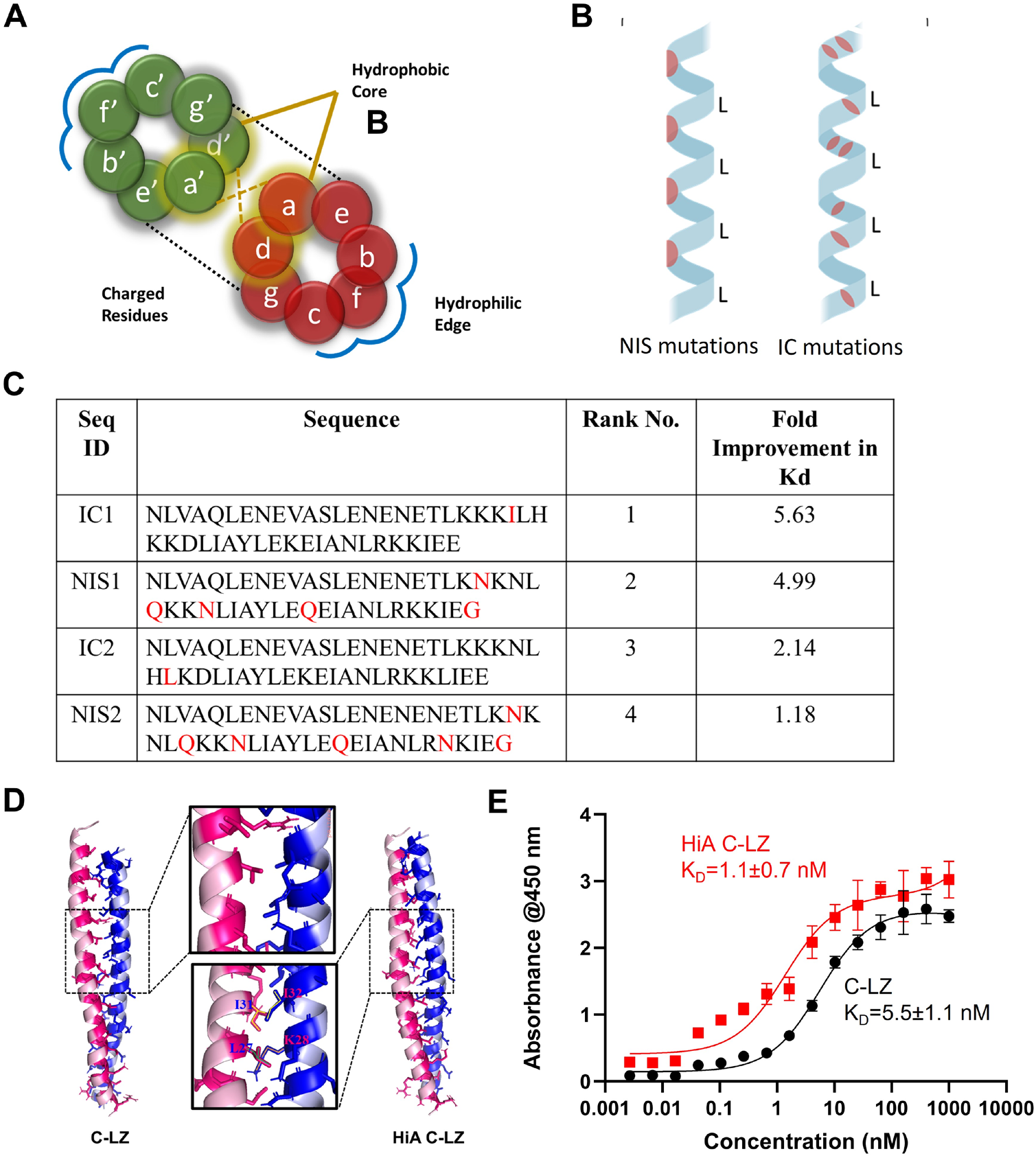
Design and characterization of high-affinity leucine zippers. (**A**) Helical wheel diagram of a leucine zipper dimer where d signifies leucine residues. (**B**) Schematic of the orientation of non-interacting surface (NIS) and interfacial contact (IC) mutations for new candidate sequences. (**C**) Fold improvement in binding affinity (Kd) of the top 4 mutant complementary leucine zipper sequences as determined by ELISA. (**D**) Representation of the increased number of intermolecular interactions of the most stable, high binding mutant protein, NIS1 (HiA C-LZ). (**E**) Binding affinity of the top HiA C-LZ (red) compared with the C-LZ as assessed by ELISA (black).

The proteins were then recombinantly expressed for in vitro testing. Using titration ELISA, the binding affinity of each sequence was tested for its affinity to the base leucine zipper (B-LZ). Upon reviewing the top four candidates (**Fig. 1C**), we selected NIS1, hereby termed high-affinity complementary leucine zipper (HiA C-LZ), for downstream studies. This decision was based on its superior binding affinity and high expression levels in E. coli. Our *in silico* results showed a significant increase in intermolecular interactions, specifically in the central region, between NIS1 and B-LZ as compared to C-LZ10nM and B-LZ (**Fig. 1D**). These findings were confirmed via ELISA, where we demonstrated that HiA C-LZ demonstrated a five-fold improvement over B-LZ binding compared with our previously tested C-LZ (**Fig. 1E**).

### Characterization of MSC Zippersomes

After optimizing leucine zipper binding affinity, we attached the HiA C-LZ to the surface of EVs to create Zippersomes [36]. This was achieved with a heterobifunctional crosslinker, Sulfo SMCC, where the amines on the EV surface were conjugated to the free thiols of either the **B-LZ** or a **C-LZ** (**Fig. 2A**). To investigate whether the improved leucine zipper binding affinity translated into increased retention and accumulation, new sequences were tested. To validate that this property could be extended to smaller vehicles, within the nanometer range, we also decorated EVs with B-LZ and C-LZ proteins to create Zippersomes. This was further validated with TEM, where unmodified EVs and scramble control Zippersomes were present as individual vesicles whereas B-LZ/HiA C-LZ Zippersomes clustered (**Fig. 2B**) with extensive contact area (**Fig. 2C**). The Zippersomes showed a 1124.23±

**Figure 2.**
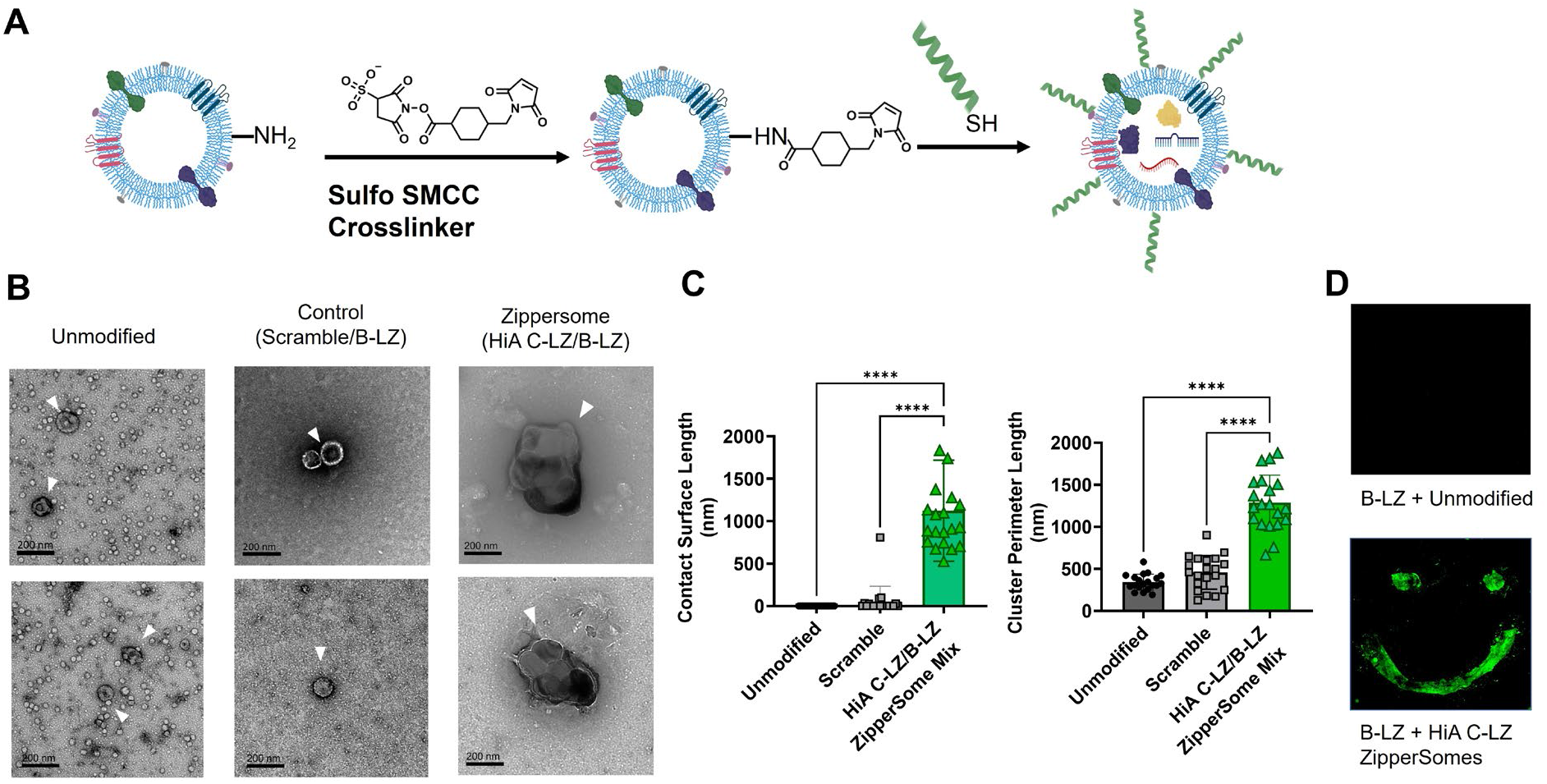
Characterization of MSC Zippersomes. (**A**) Schematic representation of cellular leucine zipper decoration using the heterobifunctional crosslinker (Sulfo SMCC), followed by maleimide-thiol conjugation. (**B**) Representative TEM images of unmodified EVs and various Zippersomes. (**C**) Calculations of contact surface length and cluster perimeter length based on TEM images (n=20). **(D)** A patterned monolayer of B-LZ incubated with fluorescently labeled unmodified EVs or HiA Zippersomes. Data are presented as mean ± SD with *p <0.05, **p<0.01, ****p<0.0001 by one-way ANOVA followed by Tukey’s post-test.

594.4 nm increase in contact surface length compared with unmodified EVs and exhibited ∼4-fold and ∼1.5-fold increases in perimeter length compared with unmodified EVs and EVs surface-decorated with a scrambled control peptide, again providing evidence of leucine zipper-mediated crosslinking. Lastly, to further validate the hetero-specificity of HiA C-LZ Zippersomes, a “smiley face” pattern of B-LZ protein was plated and blocked, and the pattern only appeared when fluorescently labeled HiA Zippersomes were incubated (**Fig. 2D, bottom row**). Without modification, fluorescently-labeled EVs were unable to bind and no detectable signal was observed (**Fig. 2D, top row**).

### Zippersomes exhibit enhanced cellular binding and uptake *in vitro*

We hypothesized that Zippersome crosslinking would enhance cellular interactions within the infarct microenvironment. To assess cellular interactions *in vitro*, four representative cell types [rat ventricular cardiomyocytes (H9c2s), fibroblasts (HCFs), endothelial cells (HUVECs), and macrophages (RAW 264.7)] were tested for EV uptake. Cardiomyocytes, fibroblasts, and endothelial cells were preconditioned in hypoxia chambers prior to EV incubation, while macrophages were treated with the hypoxia-conditioned medium (CM) along with the dosed EVs, all to effectively mimic the hypoxic infarct site (**Fig. 3A**). After three doses of alternating formulations of complementary Zippersomes, we observed that for all cell types, there was significantly more HiA Zippersome binding and uptake compared with unmodified EVs or scramble control Zippersomes (**Fig. 3B**). Additionally, H9C2 cells and macrophages bound and took up the most Zippersomes. Collectively, these results demonstrate superior cellular binding and uptake of Zippersomes compared with unmodified EVs.

**Figure 3.**
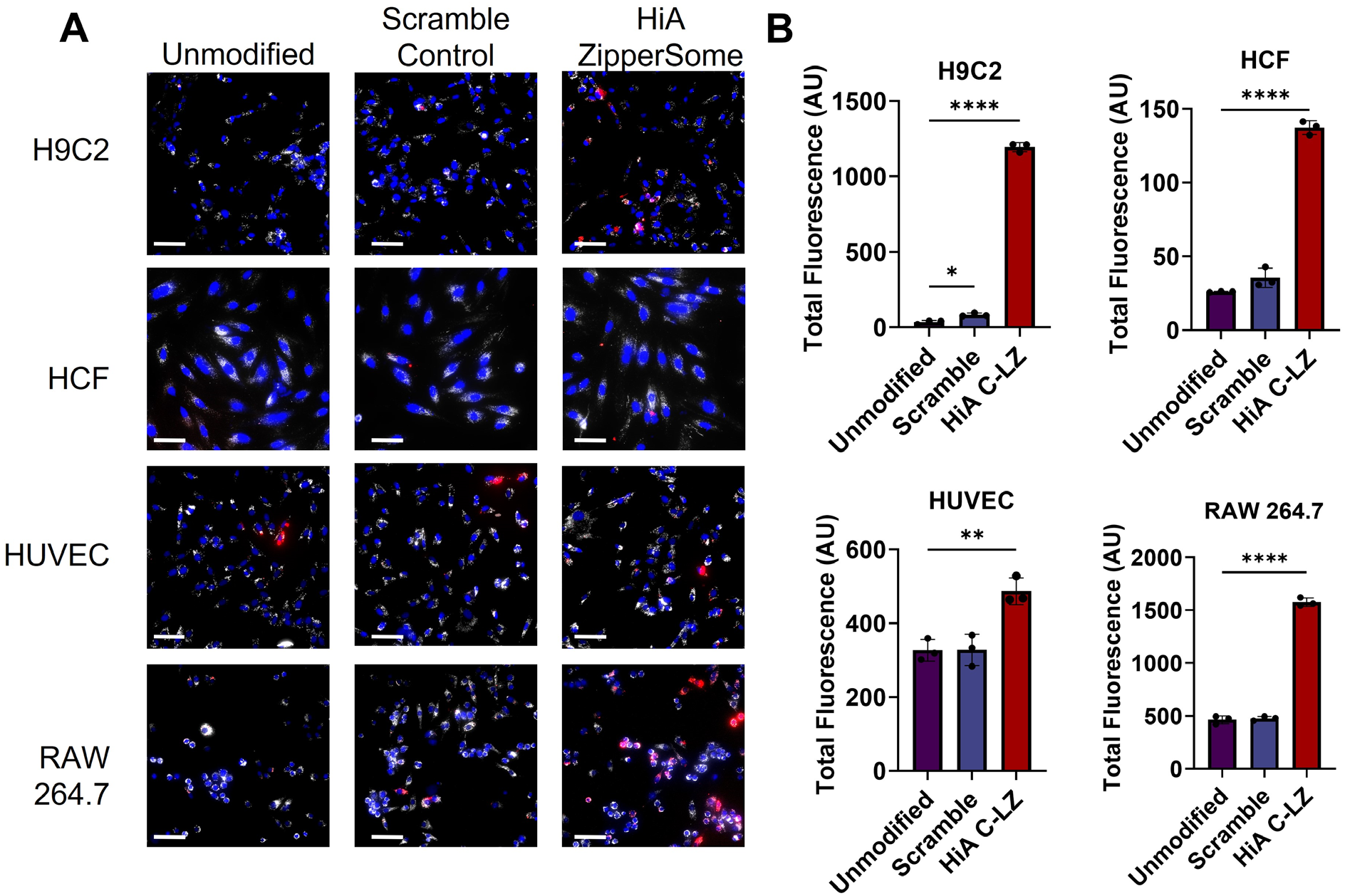
Cellular uptake of Zippersomes by cardiac cells. **(A)** Cellular uptake of DiI-labeled EVs by four representative cardiac cell types: (1) H9C2 (rat cardiomyocytes), (2) HUVECs (human umbilical vein endothelial cells), (3) HCFs (human cardiac fibroblasts), and (4) RAW 264.7 (murine macrophages). DiO was used to label cell membranes prior to incubation. Blue = DAPI, white = DiO, red = DiI. **(B)** Quantification of total fluorescence in each cell line (n=3). Data are presented as mean ± SD with *p <0.05, **p<0.01, ****p<0.0001 by one-way ANOVA followed by Tukey’s post-test. Scale bar=75 µm

### Zippersome accumulation and retention in vivo

Next, we assessed the biodistribution of DiR-labeled MSC Zippersomes after intravenous administration. EVs and Zippersomes were administered as previously described: 24 hours after MI, three alternating doses of complementary EVs were administered sequentially at 12 h intervals (**Fig. 4A**). Importantly, 60 hours post-MI, mice treated with HiA Zippersomes exhibited ∼ 7-fold greater retention than the unmodified EV control, respectively (**Fig. 4B-C**). These results show significantly improved accumulation and retention in the injured myocardium. Overall, these findings demonstrate significant shifts in biodistribution to the infarcted myocardium due to in situ leucine zipper crosslinking and persistence of our therapeutic vesicles **(Fig. 4 C, D)**.

**Figure 4.**
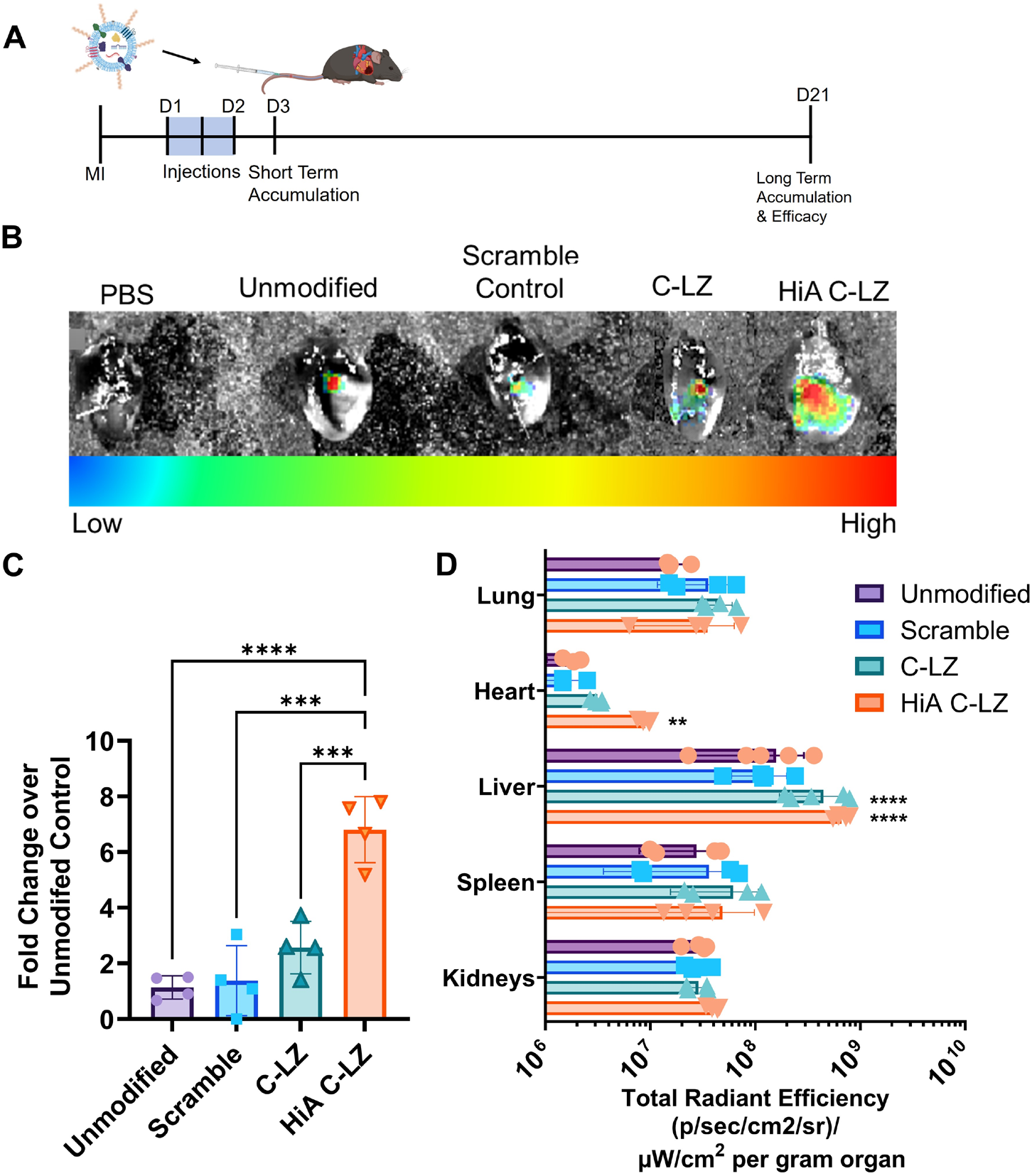
Short-term Zippersome accumulation and retention in vivo. **(A)** Schematic and experimental timeline of alternating Zippersome therapy. **(B)** Representative IVIS images using the IVIS Spectrum in vivo imaging system show DiR-labeled EV deposition in the infarcted heart, 60 hours post-MI. **(C)** Quantification of DiR fluorescence in the infarcted heart, 60 hours post-MI. **(D)** Quantification of DiR fluorescence in major organs, 60 hours post-MI. Scale bar = 50 µm. Data are presented as mean ± SD with *p <0.05, **p<0.01, ****p<0.0001 by one-way ANOVA followed by Tukey’s post-test; n=4-5.

### Zippersomes persist in the infarcted heart for up to 21 days

We next assessed if the crosslinking mediated by the leucine zippers also prolonged retention of Zippersomes in the infarcted myocardium. Excitingly, we found that there were still detectable levels of DiR signal from HiA Zippersome-treated mice within the infarcted myocardium after 21 days (**Fig. 5A**). This translated into significantly more fluorescent EV signal compared with unmodified EVs and the scramble control (**Fig. 5B**). To our knowledge, this is the longest reported retention of intravenously injected EVs in the infarcted myocardium.

**Figure 5.**
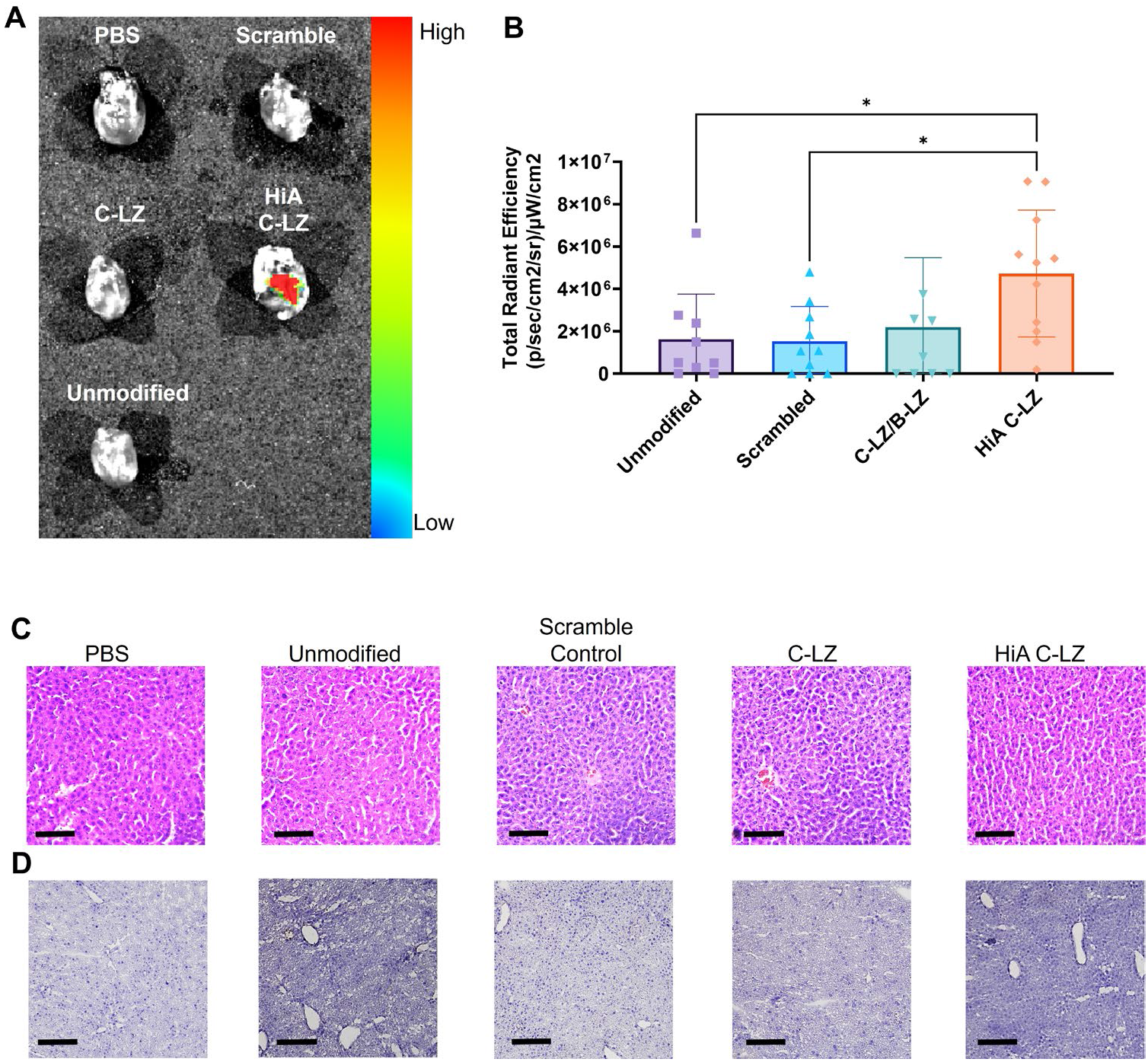
Long term EV retention in the infarcted myocardium. (**A**) Representative IVIS images of DiR-labeled EVs 21 days after the initial injection. (**B**) Quantitation of fluorescent EV signal from hearts 21 days after the initial injection. (**C**) H&E staining and (**D**) CD45 staining of liver tissues 21 days after injections. Data are presented as mean ± SD with *p <0.05, **p<0.01, ****p<0.0001 by one-way ANOVA followed by Tukey’s post-test, n=8-12. Scale bar=200µm

To evaluate the biocompatibility of the Zippersomes, we performed histopathological assessment of liver tissue on day 21 after Zippersome administration, with anti-CD45 antibodies used to identify immune cell infiltrates. There was no significant signal within any of the group, and the tissues were histopathologically normal (**Fig. 5 C, D**). These findings collectively indicate a lack of unfavorable immune responses or tissue damage in the liver.

### Zippersome treatment improves cardiac function

We next sought to determine the *in vivo* effect of intravenously injected Zippersomes on fibrosis and cardiac function after MI using a mouse model of permanent coronary artery ligation. Cardiac contractile function was tested by echocardiography in six experimental groups: sham, MI + PBS, MI + unmodified EV, MI + scramble control, MI + Zippersomes, MI + C-LZ Zippersomes, and MI + HiA Zippersomes (**Fig. 6A**). The same dosing schedule of three injections at 12 h intervals was utilized. After 21 days, HiA Zippersome-treated mice showed significantly improved cardiac function (**Fig. 6C**). HiA Zippersome-treated mice displayed significantly higher fractional shortening (FS) (33.4 ± 8.0% vs 19.6 ± 2.0%) compared with mice treated with unmodified EVs.

**Figure 6.**
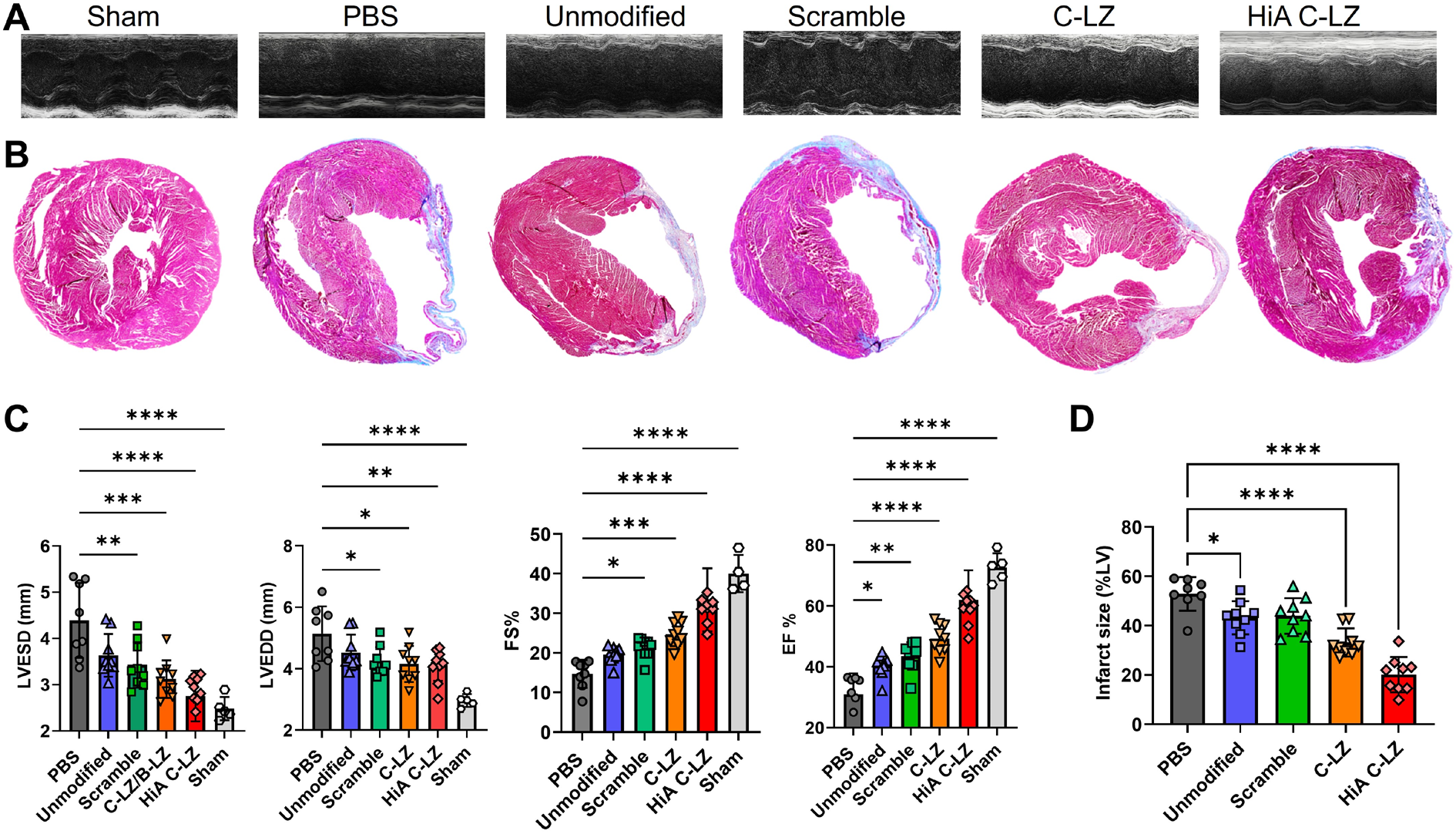
Zippersome treatment improves cardiac function and reduces infarct size. (**A**) Representative 2D M-mode echocardiography images at study completion (Day 21). Parasternal short-axis view. (**B**) Masson’s Trichrome staining was used to evaluate LV fibrotic area. (**C**) Longitudinal assessment of LV ejection fraction (EF, %), fractional shortening (FS, %), and left ventricular end diameter in systole (LVESD, mm) and diastole (LVEDD, mm). (**D**) Percentage of fibrotic LV expressed as infarct size in PBS, unmodified EV, and Zippersome hearts. Data are presented as mean ± standard deviation. Data are presented as mean ± SD with *p <0.05, **p<0.01, ****p<0.0001 by one-way ANOVA followed by Tukey’s post-test, n=5-10.

Additionally, systolic LV diameter was also substantially decreased (2.8 ± 0.6 mm vs 3.6 ± 0.8 mm), consistent with improved contractile function (**Fig. 6C**). Finally, HiA Zippersome mouse hearts had significantly less fibrosis expressed as smaller infarct size (midline length %) (20.1 ± 7.2% vs 43.2 ± 6.7%) compared with unmodified EVs (**Fig. 6B, D**). Overall, these findings demonstrate a marked improvement in multiple cardiac functional parameters after MI after HiA Zippersome treatment.

### Zippersome treatment reduces fibrosis and immune cell infiltration in spatial analyses

Finally, we aimed to examine and quantify the infiltration of immune cells such as macrophages, T cells, and B cells as well as resident myocardial endothelial cells and fibroblasts, hypothesizing that enumerating these cell populations would provide key insights into the remodeling processes occurring within Zippersome-treated hearts. To determine the spatial residencies of these model cell types, multiplex fluorescent staining and spatial analysis (**Supplemental Table 1**), paired with deconvolution, were performed. Imaris Surface creation and CytoMap software [38] were used for further analysis.

We stained for a total of 12 markers as follows: MHCII (monocytes & lymphocytes), CD11B (monocytes, granulocytes, and NK cells), CD3 (T cells), B220 (B cells), CD11C (dendritic cells), CD68 (monocytes/macrophages), CD31 (endothelial), Thy-1 (hematopoietic stem cells), Clec9a (dendritic cells), F4/80 (macrophages), Sirp α (myeloid and stem cells), and FSP (fibroblast)s. Collectively, these markers represent a host of cell types commonly used to detect cell action and residency within inflamed and wounded cite, with particular focus on detection of immune cell infiltration, fibrotic tissue, and extent of vascularization. **Figure 7** highlights fibroblasts, dendritic cells, macrophages, myeloid cells, and endogenous stem cells, which represent major influential cell types within infarcted tissues. A dynamic picture of the cell microenvironment is formed with our highly multiplexed fluorescent labeling.

**Figure 7.**
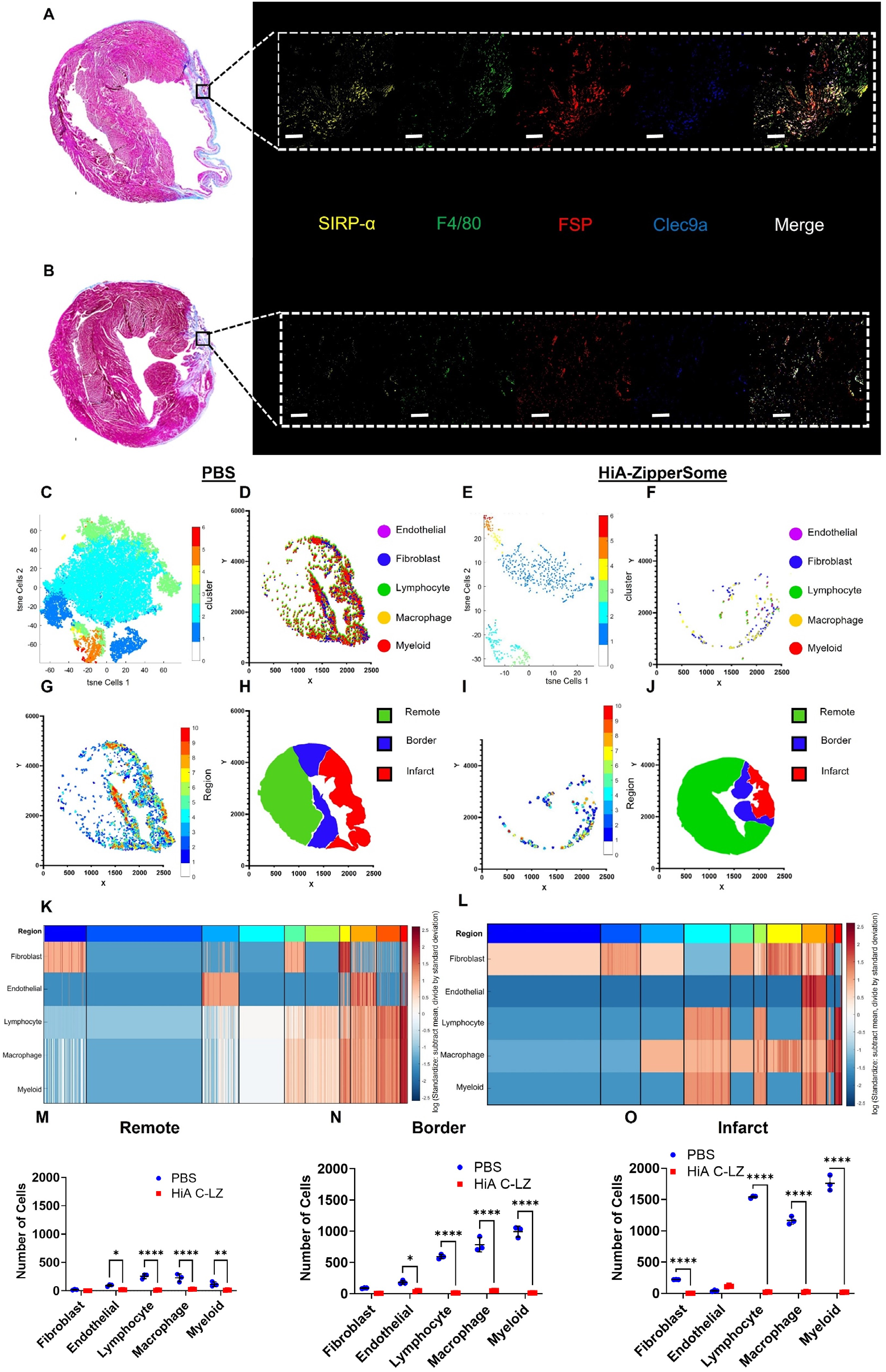
Spatial analysis of treated infarcted hearts. Representative masson trichrome and multiplex fluorescence staining of (**A**) PBS and (**B**) HiA Zippersome heart sections. (**C**) t-SNE plot of cell clusters within a representative PBS sample. (**D**) Positional plot of annotated cell clusters within a representative PBS sample. (**E**) t-SNE plot of cell clusters within a representative HiA Zippersome sample. (**F**) Positional plot of annotated cell clusters within a representative HiA Zippersome sample. (**G**) Positional plot of clustered neighborhood regions within a representative PBS sample. (**H**) Plot of myocardial cell zone surfaces within a representative PBS sample. (**I**) Positional plot of clustered neighborhood regions within a representative HiA Zippersome sample. (**J**) Plot of myocardial cell zone surfaces within a representative HiA Zippersome sample. (**K**) Heatmap of the annotated cell type clusters within a representative PBS sample. The top color bar indicates color-coded regions of neighborhoods. (**L**) Heatmap of the annotated cell type clusters within a representative HiA Zippersome sample. The top color bar indicates color-coded regions of neighborhoods. Violin plots of numbers of each cell type within the remote (**M**), border (**N**), and infarct zones (**O**) in PBS and HiA Zippersome samples. Data are presented as mean ± SD with *p <0.05, **p<0.01, ****p<0.0001 by one-way ANOVA followed by Tukey’s post-test, n=3. Scale bar=50µm.

Multiplex spatial staining captured highly complex cell populations throughout our samples (**Fig. 7A-B**). Representative fluorescence images are shown. There were significantly denser populations of detected myeloid, lymphocyte, macrophage, and fibroblast cells within PBS-treated control mouse (**Fig. 7C-H, K**) hearts than in HiA Zippersome treated mice (**Fig. 7E-J, L**). These populations, especially fibroblasts, were largely located within the infarct region and border zones corresponding to areas of fibrosis, suggesting injury-specific cell recruitment. Furthermore, HiA Zippersome hearts had significantly less cell infiltration and a more even distribution between areas.

Lastly, we assessed endothelial cells (CD31 expression) and their relative location. The percentage of endothelial cells in PBS samples was significantly smaller than those in HiA C-LZ samples (**Supplemental Figure 1**). Furthermore, endothelial cells in the PBS samples were largely localized in the border and remote areas, whereas endothelial cell signals were present in all zones of HiA Zipersome samples, including the infarct zone (**Fig 7C-J**). Further inspection revealed that within PBS mice, the majority (71.7 ± 10.8%) of identified macrophages were the proinflammatory M1 phenotype (F4/80^+^Cd11b^+^Cd11c^+^), while only a small proportion (16.9 ± 3.3%) of macrophages identified within HiA Zippersome hearts were M1 type (**Supplemental Figure 2**). Overall, these data provide insights into the primary cell types present within the injury site and potential mechanisms of action.

## Discussion

MSC-derived EVs have been used as a promising alternative to live cell therapies for the treatment of MI and wound healing[40–44]. Many studies have now shown that EVs are key mediators of stem cell-mediated wound healing and regeneration within the damaged heart, largely due to their ability to move, even through occluded sites. However, despite their widespread use within the field, they are often implemented via direct myocardial injection, which poses risks and does not overcome limited retention [43,45,46]. In addition to the inherent risk and limited translatability of this invasive administration route, rapid biodistribution and clearance of these vehicles is often observed, so the therapeutic effects are often limited at the target site. The Zippersomes introduced in this study are a novel method to address these hurdles.

Here we aimed to reduce invasive procedures, improve cellular uptake, and accumulate EVs at the infarct site for an extended period to improve therapeutic efficacy and cardiac function long-term. We hypothesized that leucine zipper crosslinking could also be used to enhance the accumulation and retention of EVs to increase cellular uptake and therefore improve cardiac function. To facilitate crosslinking of EVs, we attached heterodimerizing leucine zippers to two sets of EVs[47,48], which were subsequently dosed in an alternating manner. We previously showed that the leucine zippers are biocompatible and can stably bind long term within serum conditions to facilitate cellular crosslinking in vivo[19].

The cellular uptake of EVs is important to consider, since their circulatory half-life is typically very brief (t1/2 < 30 min [49,50]). By increasing the number of EVs clustering at the cell surface, we hypothesized that overall cardiac retention and cellular uptake could be enhanced due to stabilizing effects. Increased cellular uptake of EVs is particularly important within inflammatory disease sites, such as infarct sites, where EV persistence is necessary to regulate inflammatory responses [51–53][54,55]. First, we found that when complementary Zippersome populations are mixed, they can crosslink and form large, defined clusters. Furthermore, we found that crosslinking and cluster formation increased cellular binding and uptake, especially within cardiomyocytes and macrophages. Previous studies have shown that MSC-derived EVs possess anti-inflammatory effects and are able to polarize inflammatory macrophages present in the infarcted myocardium after MI, towards an anti-inflammatory phenotype, decreasing the number of inflammatory M1 macrophages[56–58], suggesting a possible mechanism of action for EV-mediated regeneration in vivo.

Unmodified EVs are typically cleared within hours of local or systemic administration. Encouragingly, we found that through our vehicle surface decoration, we increased cardiac retention of MSC EVs up to day 21. To our knowledge, this level of retention, particularly through systemic administration, has never been demonstrated before: published studies typically report retention of up to 24 hours [59]. Although we also observed slightly enhanced retention of Zippersomes within livers compared with unmodified EVs, we did not observe signs of toxicity or immune infiltrations within these livers.

Additionally, we discovered that Zippersomes, especially those containing the high binding affinity leucine zipper (HiA C-LZ), enhanced the therapeutic effect of MSC EVs and substantially improved cardiac function, likely due to increased retention at the target site. On day three, we observed a ∼ 7-fold increase in Zippersome accumulation within the heart compared with unmodified EVs. On day 21, we found that cardiac contractile function substantially increased while infarct size and LV diameter significantly decreased compared with both PBS- and unmodified EV-treated mice. Notably, the Zippersomes were able to restore cardiac function of mice back to healthy levels. While most other EV MI studies have shown modest improvements in fibrosis in the range of approximately 10-20% compared with vehicle-treated control animals [42,60–62], we observed an impressive 60% improvement in infarct size reduction. Collectively, these findings support the hypothesis that retention can be controlled by altering the leucine zipper binding affinity.

Upon spatial cell analysis, PBS samples showed extensive fibroblast and immune cell residency within sites of injury, consistent with well-characterized post-infarct pathobiology[63–67]. We detected the presence of dense mixtures of lymphocytes and myeloid cells, interspersed with pockets of fibroblasts. Previous studies have shown that myeloid cells and lymphocytes infiltrate ischemic and infarcted areas, where they work in tandem as part of the collective immune response to remodel the myocardium[68,69]. Myeloid cells typically infiltrate first, as part of the innate immune response, followed by lymphocytes as part of the adaptive immune response. Using spatial mutiomic analysis, Kuppe et al. also discovered strong dependencies between lymphocytes and myeloid cells, which is supported by our observed colocalization of these cell types, largely within the infarct and border zones of PBS treated mice [70]. Studies have shown that excessive accumulation of these cells within the heart following an injury can also increase reactive oxygen species and amplify fibrotic injury[71], which may exacerbate fibrosis.

Nahrendorf and colleagues showed that excessive infiltration of inflammatory macrophages can suppress cardiac repair through various pathways such as dysregulated extracellular matrix remodeling and altered cytokine expression. Therefore, inhibiting inflammatory macrophage infiltration and increasing the relative proportion of anti-inflammatory macrophages can improve cardiac function and decrease infarct size[72–76]. This mechanism is supported by our results where we found that within the HiA ZipperSome-treated mice, there was a substantial reduction in the detection of all inflammatory cell types, including macrophages, likely indicating a reduced inflammatory environment, resulting in significant reduction in fibrosis. Furthermore, within the PBS-treated control mice, >70% of macrophages were the pro-inflammatory M1 phenotype (F4/80^+^Cd11b^+^Cd11c^+^), while the number of M1 macrophages identified within HiA Zippersome hearts substantially decreased to ∼16.9% (**Supplemental Figure 2**). Ultimately this points to inflammatory macrophage reduction and macrophage polarization as another possible mechanism of action for our observed ZipperSome mediated cardiac repair.

One major advantage of our system is its high level of tunability. In the future, our zipper-based crosslinking technology could be combined with various therapeutic cargoes and drug carriers. Additionally, Zippersomes could be applied to different disease states where enhanced retention could provide therapeutic benefits where targetable surface areas are limited. Because each therapeutic dose is designed to capture the next therapeutic dose, we have developed a highly effective strategy to increase drug accumulation and retention. Intravenous injection was selected as the route of administration due to its clinical translatability. Typically, patients diagnosed with MI receive continuous IV infusions throughout their hospital stay, making it a convenient and accessible route for multiple dosing strategies.

## Conclusion

In conclusion, through simple surface engineering, leucine zippers can be used to generate EVs capable of amplifying the targetable surface areas and crosslinking in situ. We demonstrate that sequence-specific crosslinking improves cellular uptake for more potent therapeutic mechanisms. We show that intravenously administered Zippersomes accumulate in the injured myocardium and are retained for several weeks. Long-term retention significantly improved cardiac function, as shown by a remarkable decrease in fibrosis, and an improvement in ejection fraction and fractional shortening. This novel method provides proof of concept for significant improvement in EV delivery and other drug delivery carriers.

## Acknowledgments

We acknowledge funding through the National Institutes of Health (1R01HL161456, R01GM150252, R01CA241679), the Eshelman Institute of Innovation, and the National Science Foundation (DMR 2000256). JN is an inventor on a patent filing related to the compositions and methods for programming extracellular vesicles, which is licensed by ExoPharm. However, that technology is not part of this paper. Parts of figures 2 and 3 were created using Biorender.com. Confocal images were acquired at the UNC Neuroscience Microscopy Core 593 (RRID:SCR_019060), which is supported, in part, by funding from the NIH-NINDS 594 Neuroscience Center Support Grant P30 NS045892 and the NIH-NICHD Intellectual and Developmental Disabilities Research Center Support Grant P50 HD103573. IVIS images were acquired at the UNC Biomedical Research Center. We also acknowledge funding from the Pharmaceutical Research and Manufacturers of America (PhRMA) Foundation for N.Jasiewicz.

## Data Availability Statement

The data that support the findings of this study are available from the corresponding author, JN, upon reasonable request.

## Code Availability Statement

All code used for analysis is available at https://gitlab.com/gernerlab/cytomap

## Sample Statement

All measurements were taken from distinct samples.

## Methods & Materials

### Molecular docking & in silico binding

Three-dimensional structural models of all leucine zipper proteins were generated using ITASSER [26,27] based on the known structure of homologous proteins. Energy-minimized models of base leucine zippers (B-LZ) and complementary leucine zippers (C-LZs) were molecularly docked using HADDOCK [28,29] to generate dimers.

Amino acids designated within the “A, D, E, and G” heptad positions were marked as active residues, and a single model was chosen as the final conformation. Binding affinity and intermolecular interactions (Kd) were determined using PRODIGY[30].

### Protein expression & purification

All leucine zipper sequences were cloned into a pET21a vector (Novagen, Burlington, MA) and expressed in BL-21 (DE3) cells (Lucigen, Middleton, WI). Bacterial cultures were grown overnight at 37°C in 2x YT (Fisher Scientific, Waltham, MA) medium and ampicillin. For large-scale production, a 10 L autoinduction[31] culture was inoculated with 1% of the overnight culture and grown for four hours at 37°C and then an additional 20 hours at room temperature. After incubation, cells were harvested 16 h post-induction by centrifugation. The cell pellet was resuspended in lysis buffer containing 50 mM Tris, 100 mM KCl, 10% glycerol, 100 µg/ml lysozyme, and 1 mM Phenylmethanesulfonylfluoride Fluoride (PMSF), pH 8.0. The cell suspension was sonicated and then centrifuged at 17,000 x g for 40 minutes at 4°C, after which Ni-NTA Resin (G-Biosciences, St. Louis, MO) was added to the solution. The beads were incubated with the solution and the protein was isolated by immobilized metal affinity chromatography (IMAC)[32–34]. Endotoxin removal was performed using Pierce High-Capacity Endotoxin Removal resin (Thermofisher) and validated using the Pierce LAL Chromogenic Endotoxin Kit Quantitation Kit (Thermofisher).

### Enzyme-linked immunosorbent assay (ELISA)

Nunc MaxiSorp 96-well plates were coated with Base-Leucine Zipper (B-LZ) at 2.5 µg/mL and incubated overnight at 4°C. The plate was thoroughly washed with 0.1% TBS-Tween (TBST). The wells were washed with 0.1% PBST and then blocked with BSA blocking buffer for 1 h at room temperature before incubation with complementary leucine zipper (C-LZ) concentrations ranging from 1 µM to 0.3 µM at room temperature for 2 hours before washing. Washes were followed by the addition of 100 μL of anti-FLAG horseradish peroxidase (HRP)-linked monoclonal antibody (1:10,000 mAb, #A8592, Sigma Aldrich, St. Louis, MO) for 1 h at room temperature with rocking. The wells were washed with 0.1% PBST followed by the addition of TMB-ELISA substrate. After a 10 min incubation, 2 M H2SO4 was added to stop the reaction. The absorbance was measured at 450 nm and 570 nm with a SpectraMax M5 plate reader (Molecular Devices, Sunnyvale, CA)[35].

### Cell culture and EV isolation

Conditioned medium was collected from murine bone-marrow-derived mesenchymal stem cells (Cyagen MUBMX-01101) grown to 70% confluency and then incubated with EV-depleted FBS-medium for 48 hours. EVs were isolated by differential centrifugation according to previously established protocols[36,37].

### Protein-EV conjugation

EV concentration was determined by nanoparticle tracking analysis (NTA). At a concentration of 1.0e10 particles/ml, cells were incubated with 2.5 µM DiR [DiIC18(7) (1,1’-dioctadecyl-3,3,3’,3’-tetramethylindotricarbocyanine iodide] dye and incubated with 500 µg sulfosuccinimidyl 4-(N-maleimidomethyl) cyclohexane-1-carboxylate (Sulfo-SMCC) for 30 minutes at 37°C. To reduce protein for conjugation, proteins were incubated with 1 mM tris(2-carboxyethyl)phosphine (TCEP) at room temperature for 20 minutes. After washing EVs with Amicon Ultra-15 100 kDa MWCO centrifugal filter units, 2 mg of protein was added to EVs and incubated at room temperature for 30 minutes before final washing and resuspension.

### EV characterization

EVs were prepared as previously described[36], and their concentration and size were evaluated by NTA (Particle Metrix, ZetaView). For transmission electron microscopy (TEM), isolated EVs were incubated with glow discharged Formvar/carbon-supported copper TEM grids. Samples were allowed to adsorb on the grid for 2 min before wicking off with a filter paper. 1% uranyl acetate was applied to the grid for staining, and the excess was wicked off immediately and repeated again. Finally, grids were allowed to air dry and then imaged using the FEI Tecnai T12 TEM at 120 V [36].

### Enzyme-linked immunosorbent assay (ELISA)

Nunc MaxiSorp 96-well plates were coated with leucine zippers at 2.5 µg/mL and incubated overnight at 4°C. The plate was thoroughly washed with 0.1% TBS-Tween (TBST). Wells were washed with 0.1% PBST and then blocked with BSA blocking buffer for 1 h at room temperature before incubation with a stock of 3.3 μM C-LZ at 37°C for 2 hours before washing. Washes were followed by the addition of 100 μL of anti-FLAG horseradish peroxidase (HRP)-linked monoclonal antibody (mAb, #A8592, Sigma Aldrich, St. Louis, MO) for 1 h at room temperature with rocking. The wells were washed with 0.1% PBST followed by the addition of TMB-ELISA substrate. After 10 minutes of incubation, 2 M H2SO4 was added to stop the reaction. The absorbance was measured at 450 nm and 570 nm with a SpectraMax M5 plate reader (Molecular Devices, Sunnyvale, CA).

### Patterned EV binding

Individual glass microscope slide sections were circled with a hydrophobic pen and were coated with poly-D-lysine (0.01% solution) for 1 hour at room temperature. After washing three times with deionized water, the slides were wrapped tightly in a single layer of parafilm. Patterns were cut into the parafilm using a 30-gauge needle and then incubated with B-LZ (2.5 mg/ml) solution for three, 1 h incubations. The parafilm was removed and the outlined regions were blocked with 5% milk solution for 1 hr. Sections were then incubated with DiI (DiIC18(3); 1,1′-dioctadecyl-3,3,3′,3′-tetramethylindocarbocyanine) labeled unmodified EVs or Zippersomes at a concentration of 1e14 particles/ml for three hours, followed by washing, and mounting with Fluorsave (Millipore, Burlington, MA). Images were taken with the Nikon Eclipse Ti2 microscope.

### Cellular EV Uptake

Unmodified EVs and Zippersomes, were labeled with DiI (1,10-dioctadecyl-3,3,30,30-tetramethylindocarbocyanine perchlorate). RAW 264.7 macrophages, human umbilical vein endothelial cells (HUVECs) (ATCC, Cat# CRL-1730), HL-1 cardiomyocytes (Millipore, Cat# SCC065), and human cardiac fibroblasts (HCFs) (Sciencell, Carlsbad, CA; Cat# 6300) cells were used as representative cell types within the infarct microenvironment. Cells were seeded at a density of 5,000 cells/well in 96 well plates and left to adhere overnight. HUVECs, HL-1s, and HCFs were maintained in hypoxic conditions (5% CO2, 5% O2) for 24 hours prior to studies. After 24 h, conditioned media (CM) from all cell types were collected, mixed, and dosed to RAW 264.7 cells to mimic infarct microenvironment exposure. The remaining cells were given fresh medium. Cells were treated with doses of 1.11e8 EVs/well each hour, followed by washing for a total of three doses within the first three hours. Eight hours after the first dose, wells were washed and fixed with 4% paraformaldehyde before imaging with a Nikon Eclipse Ti2 microscope.

### Animals

Female 10–12-week-old Female C57BL/6 mice were used for the in vivo studies. The University of North Carolina at Chapel Hill Institutional Animal Care and Use Committee (IACUC) approved all animal procedures. All methods and experiments were performed in accordance with the U.S National Institutes of Health Guide for Care and Use of Laboratory Animals. Humane care and treatment of animals were ensured.

### Mouse model of MI (permanent coronary ligation)

All surgeries were performed in the McAllister Heart Institute (MHI) Cardiovascular Physiology and Phenotyping Core. Briefly, mice were anesthetized with isoflurane. An incision was made to visualize the trachea before intubating with a 20-gauge blunt needle and ventilation. A left lateral thoracotomy exposed the heart, and the left coronary artery (LCA) was identified and permanently occluded with a 7-0 nylon suture. Occlusion was confirmed by electrocardiography (ECG) and visual inspection of myocardial pallor. The thorax was closed in layers (ribs, muscles, and skin). Mice were provided with analgesics and monitored per protocol.

### Biodistribution of EVs

For biodistribution studies, 1.3e9 DiR-labeled Zippersomes or unmodified MSC derived EVs, or PBS was administered via tail vein injection into C57BL/6 mice 24 hours after inducing MI. A total of three injections were administered every twelve hours. Twelve hours after the final injection (60 hours post-MI or 21 days post-MI), mice were sacrificed and the brain, lung, heart, liver, spleen, and kidneys were collected and weighed.

Fluorescent biodistribution was analyzed using the IVIS Spectrum in vivo imaging system (PerkinElmer, Waltham, MA). Average region of interest (ROI) signals were calculated using Aura Imaging Software (Spectral Instruments, Tucson, AZ). The data is presented as total radiant efficiency/g organ.

### Histopathological assessment

For histolopathological studies, 1.3e9 DiR-labeled Zippersomes or unmodified MSC EVs, or PBS were administered via tail vein injection into C57BL/6 mice. A total of three injections were administered every twelve hours.

1. *Assessment of infarct size*: 21 days post-MI, mice (n=8-10) were briefly perfused with KCl (30 mM) to arrest the heart in diastole, sacrificed, and hearts collected. Harvested hearts were fixed in formalin for two days, dehydrated, and then cleared before paraffin embedding. Tissues were then cut into 7.5 μm sections, which were deparaffinized and rehydrated for subsequent staining. Masson’s trichrome staining was performed using a kit (Sigma-Aldrich, Burlington, MA). Whole heart images were taken with the Nikon Eclipse Ti2 microscope. Midline left ventricle lengths were measured using ImageJ software.
2. Histological assessment of CD45 signal in livers: After day 21 sacrifice, livers were frozen and embedded in optimal cutting temperature (OCT) compound. Tissues (n=3) were cut as 8 µm sections and immediately fixed in ethanol. Sections were then incubated with anti-mouse CD45 (Tonbo Biosciences, San Diego, CA; Cat # 70-0451-U100) at a 1:200 dilution for 30 minutes at room temperature. This was followed by an HRP secondary antibody (Tonbo Biosciences, San Diego, CA, Cat # 72-8104-M001). Chromogen was developed with a DAB HRP substrate (Vector Labs, Newark, CA; Cat # SK-4105). Samples were then counterstained with hematoxylin and bluing reagent (Biovision, Milpitas, CA) before mounting with Permount. Images were taken with the Nikon Eclipse Ti2 microscope.

### Echocardiography

Echocardiograms were collected with the Vevo2100 Ultrasound system (VisualSonics, Toronto, Canada). Two-dimensional M mode parasternal short axis views were recorded. Left ventricular dimensions, wall thickness, heart rate, and cardiac output were recorded. Left ventricle (LV) stroke volume (SV) was calculated as the difference between the end diastolic volume (EDV) and the end systolic volume (ESV). Ejection fraction (EF) was calculated as (EF = [(EDVESV)/EDV] *100).

### Spatial analysis

Serial sections from histological tissue assessment were utilized for fluorescence staining (7.5 µm cleared paraffin-embedded sections, prepared as previously described). Sections were stained with a panel of 12 fluorescently conjugated antibodies, shown in Table S1. Next sections were cover slipped and mounted with Fluorsave (Millipore) and imaged using mosaic tile imaging on an Olympus FV3000RS confocal microscope.

For image analysis, tiles were exposed to flatfield correction (BioVoxxel, ImageJ) and then deconvolved and stitched using Huygens software (Scientific Volume). Stitched files were imported into Imaris, and downstream analysis was performed as outlined in the CytoMap Spatial Analysis methods [38].

### Statistical analysis

All the quantitative data were presented as mean ± SD. The mean values are based on a minimum of n=3 biological replicates. Groups were compared using one-way ANOVA with a post hoc Tukey test (*p<0.05, **p<0.01, ***p<0.001). All statistical analyses were performed using Prism 7 (GraphPad Software, La Jolla, CA). The error bars represent standard deviations (SD).

