## Supplemental Information for "In Situ-Crosslinked Zippersomes Enhance Cardiac Repair by Increasing Accumulation and Retention"

| Ch | Antibody | Fluorophore |
|---|---|---|
| 1 | MHCII | biotin (+ Streptavidin-v500) |
| 2 | CD11B | BV605 |
| 3 | CD3 | BUV395 |
| 4 | B220 | spark violet 538 |
| 5 | CD11C | BV480 |
| 6 | CD68 | VioBlue |
| 7 | CD31 | BV421 |
| 8 | THY1 | AF488 |
| 9 | CLEC9A | PE |
| 10 | F4/80 | APC |
| 11 | SIRP | AF594 |
| 12 | FSP | CF633 |

**Supplemental Table 1:** Antibody staining panel for spatial analysis. Antibodies and their conjugated fluorophore used for multiplex spatial analysis of ZipperSome and PBS treated mouse hearts.

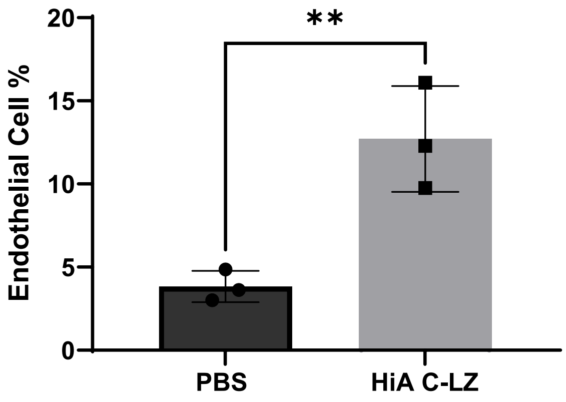


**Supplemental Figure 1: Endothelial cell percentage of total cell count from spatial analysis.** Comparison of percentages of endothelial cells out of total cells detected in PBS-treated mice and HiA-ZipperSome treated mice 21 days after myocardial infarction via spatial analysis. Data are presented as mean ± SD with *p <0.05, **p<0.01, ****p<0.0001 by unpaired t test. N=3

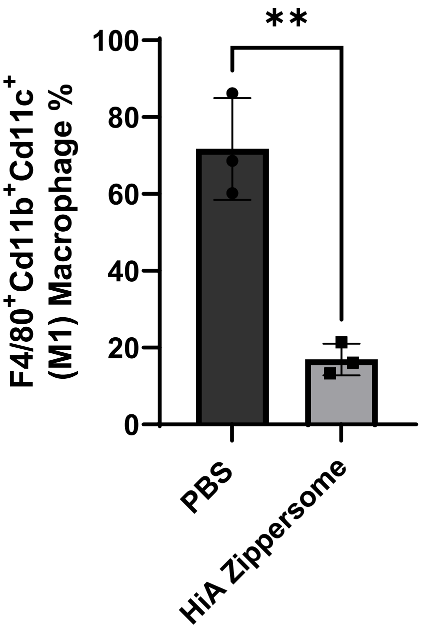


**Supplemental Figure 2: M1 macrophage composition**. Comparison of percentages of pro-inflammatory (M1) phenotypic cells out of total detected macrophages within heart samples of PBS treated mice and HiA-ZipperSome treated mice. Data are presented as mean ± SD with *p <0.05, **p<0.01, ****p<0.0001 by unpaired t test. N=3.
